# Fork Pausing Complex Engages Topoisomerases at the Replisome

**DOI:** 10.1101/738328

**Authors:** Maksym Shyian, Benjamin Albert, Andreja Moset Zupan, Vitalii Ivanitsa, Gabriel Charbonnet, Daniel Dilg, David Shore

## Abstract

Replication forks temporarily or terminally pause at hundreds of hard-to-replicate regions around the genome. A conserved pair of budding yeast replisome components Tof1-Csm3 (fission yeast Swi1-Swi3 and human TIMELESS-TIPIN) acts as a ‘molecular brake’ and promotes fork slowdown at proteinaceous replication fork barriers (RFBs), while the accessory helicase Rrm3 assists the replisome in removing protein obstacles. Here we show that Tof1-Csm3 complex promotes fork pausing independently of Rrm3 helicase by recruiting topoisomerase I (Top1) to the replisome. Topoisomerase II (Top2) partially compensates for the pausing decrease in cells when Top1 is lost from the replisome. The C-terminus of Tof1 is specifically required for Top1 recruitment to the replisome and fork pausing but not for DNA replication checkpoint (DRC) activation. We propose that forks pause at proteinaceous RFBs through a ‘sTOP’ mechanism (‘slowing down with TOPoisomerases I-II’), which we show also contributes to protecting cells from topoisomerase-blocking agents.

## INTRODUCTION

The chromosomal DNA of most cells is duplicated once per cell cycle due to the concerted action of DNA helicases unwinding the DNA template, topoisomerases unlinking the parental strands, and DNA polymerases synthesizing the daughter strands in collaboration with a myriad of accessory factors (Bell and Labib 2016). This assembly of proteins on the DNA replication fork is called the ‘replisome’. In order to achieve the completeness of genome duplication, the replisome should pass through the entirety of all chromosomes. On average budding yeast replisomes move through ca. 20 kb of DNA before merging with a converging fork (Pasero et al. 2002). However, *in vivo* the speed of the replisome is not uniform, as it temporarily or terminally slows/pauses/arrests/stalls at certain locations, called replication fork barriers (RFBs). RFBs are comprised by ‘unconventional’ DNA structures (inverted repeats, trinucleotide repeats, G4 quadruplexes), RNA/DNA hybrids (R-loops), and tight protein/DNA complexes (Gadaleta and Noguchi 2017). Examples in yeast of the latter type of RFB are found at the rDNA repeat array, tRNA genes (tDNA), telomeres, centromeres, silent mating type loci (*HML*/*HMR*) silencer elements, and dormant origins of replication (Gadaleta and Noguchi 2017).

Replisome pausing at these protein barriers involves two components: (1) a tight DNA-binding protein block specific for a given locus (e.g. Fob1 (rDNA RFB - rRFB), the RNA polymerase III pre-initiation complex, the general regulatory factor Rap1, or the origin recognition complex) and (2) a “fork pausing/protection complex” (FPC) – the evolutionary conserved heterodimer represented by Tof1-Csm3 in budding yeast (Swi1-Swi3 in fission yeast and TIMELESS-TIPIN in human). Tof1-Csm3 is also found in association with Mrc1 (not itself involved in replication pausing) in a trimeric complex referred to as MTC, which travels with other factors in a still larger assembly on replication forks called the Replisome Progression Complex (RPC) (Gambus et al. 2006). Loss of Tof1-Csm3 leads to a decrease in replisome pausing at many of the studied protein barriers in budding and fission yeast, and human cells (Gadaleta and Noguchi 2017), while increasing blockage at some unconventional DNA structures (Voineagu et al. 2008). Accessory 5’ to 3’ DNA helicase Rrm3 is a part of the yeast replisome and uses its ATPase/helicase activity to assist the main replicative 3’ to 5’ CMG helicase (Cdc45-Mcm2-7-GINS) in progression specifically at the protein blocks (Ivessa et al. 2000; Ivessa et al. 2003; Azvolinsky et al. 2006). Replication fork stalling is proposed to fuel tumorigenesis and ageing (Gaillard et al. 2015). However, the molecular mechanism of action of the Tof1-Csm3, Rrm3 and replisome progression through protein blocks is complex and incompletely understood.

In addition to helicases, the replisome must employ topoisomerases in order to topologically unlink, or swivel, the two parental DNA strands (Duguet 1997). Topoisomerase I (Top1 in budding yeast) is regarded as the main replicative swivelase, while topoisomerase II (yeast Top2) provides a back-up mechanism when Top1 is not available (Kim and Wang 1989a; Bermejo et al. 2007). It was postulated that similarly to helicases, topoisomerase action should be impeded by the presence of tight protein complexes on DNA in front of the fork (Keszthelyi et al. 2016).

We set out here to understand the mechanism of Tof1-Csm3-dependent replisome arrest/pausing at RFBs. We show first that the Tof1-Csm3 fork pausing complex acts independently of the accessory helicase Rrm3. Instead, we find that Tof1-Csm3 engages replicative topoisomerase I (and backup topoisomerase II) at the replisome to promote fork pausing at proteinaceous RFBs (sTOP mechanism). The Tof1 C-terminus mediates Top1 association with the replisome and fork pausing but is not required for the DNA replication checkpoint (DRC). sTOP and DRC mechanisms jointly promote cellular resistance to topoisomerase-blocking agents.

## RESULTS

### Fork pausing complex Tof1-Csm3 acts independently of Rrm3 helicase

Replication forks slow down at hundreds of tight protein/DNA complexes around the yeast genome (Gadaleta and Noguchi 2017). In search for the fork pausing mechanism, we started by first confirming with 2D and 1D gels (Brewer and Fangman 1988; Kobayashi et al. 2004) that only Tof1-Csm3 but not Mrc1 (Tourriere et al. 2005; Hodgson et al. 2007) or other related RPC components are required for fork pausing at the rRFB (**Fig. 1A, S1A-C**). Accessory helicase Rrm3 helps the replisome to move past protein RFBs throughout the genome (Ivessa et al. 2003). Upon initial characterization of the roles of Tof1 and Csm3 in fork pausing using 2D gels, it was postulated that they work by counteracting the Rrm3 helicase (**Fig. 1A**, model ‘1’) (Mohanty et al. 2006; Bairwa et al. 2011). If this were true, fork pausing should become completely independent of Tof1-Csm3 in cells lacking Rrm3. However, closer inspection of the 2D gel evidence in the above initial reports suggests that this was not the case.

**Figure 1.**
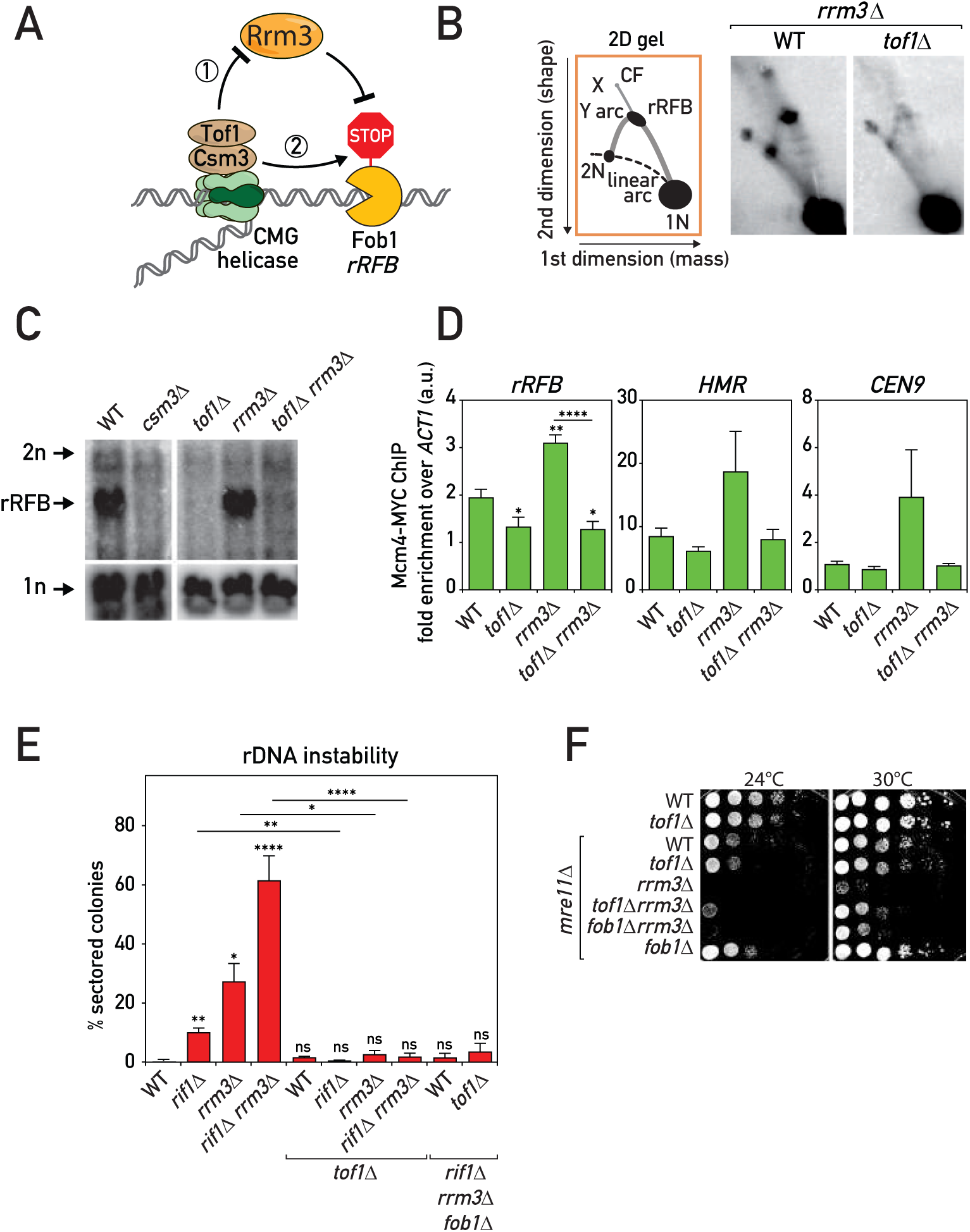
Tof1-Csm3 complex functions independently of Rrm3 helicase (A) Schematics of Rrm3-dependent (‘1’) and -independent (‘2’) mechanisms for Tof1-Csm3 role in replication fork pausing at proteinaceous barriers. (B-D) *tof1*Δ suppressed fork pausing in *rrm3*Δ cells: (B) Schematic (left) and representative images (right) of replication intermediates detected in the asynchronous cultures of strains of indicated genotypes by Southern hybridization with rDNA rRFB probe on *Bgl* II-digested DNA separated with 2D gels and blotted to nylon membrane; (C) same as in (B) but Southern blot done directly on 1st dimension gels; (D) Replisome pausing detection with Mcm4-MYC ChIP-qPCR at several pausing sites in asynchronous cultures of strains of the designated genotypes. (E) *tof1*Δ suppressed rDNA instability in *rrm3*Δ and *rif1*Δ cells – rDNA instability measurement with *ADE2* marker loss assay. (F) *tof1*Δ partially alleviated *mre11*Δ *rrm3*Δ synthetic sickness – serial dilution growth assay. X – X-shaped molecules; CF – converging forks. Means with SEM are plotted; Welch’s t test was used for quantitative comparisons (*p<0.05; **p<0.01; ***p<0.001; ****p<0.0001; ns – not significant). See also Fig. S1.

To clarify the Tof1-Csm3 relationship with Rrm3 we utilized several replication fork pausing and instability assays (**Fig. 1**). Deletion of *TOF1* or *CSM3* led to a strong decrease in paused fork signal at Fob1-RFB detected by 1D gels, as expected (**Fig. 1C**). Significantly, *tof1Δ* mutation also decreased fork pausing in a *rrm3Δ* background (**Fig. 1B, 1C**), suggesting that in cells lacking Rrm3 helicase, Tof1 still actively promotes replication fork slowdown (**Fig. 1A**, model ‘2’). Next, we used chromatin immunoprecipitation to probe binding of the replicative helicase components Mcm4 and Cdc45, as it was reported that replisome components are more enriched at pause sites (Azvolinsky et al. 2009). Consistent with the 1D and 2D gel analysis, we detected Tof1-dependent enrichment of Mcm4 and Cdc45 on several pause sites in cells lacking Rrm3 helicase (**Fig. 1D** and **S1D**), while pausing at telomeres was less dependent on Tof1.

Lack of the Rrm3 helicase leads to prolonged fork pausing at Fob1-RFB and elevated rDNA instability as a result of fork pausing (Ivessa et al. 2000). Utilizing *ADE2* marker loss from the rDNA locus as a measure of ribosomal gene array instability, we found that Tof1 was required for rDNA repeat destabilization in *rrm3Δ* cells (**Fig. 1E**). Remarkably, *tof1Δ* mutation also suppressed the more elevated instability of an *rrm3Δ rif1Δ* double mutant, which additionally lacks a negative regulator of replication origin firing, Rif1 (Shyian et al. 2016). Viability of *rif1Δ* cells requires the DSB repair and fork maintenance complex MRX, and the lethality caused by MRX mutations in these cells is suppressed by pausing alleviation through *fob1Δ*, *tof1Δ*, or *csm3Δ* mutations (Shyian et al. 2016). Notably, we observed that *tof1Δ* partially suppressed synthetic sickness of *rrm3Δ* and *mre11Δ* mutations, to an extent slightly stronger than suppression by *fob1Δ* (**Fig. 1F**). This difference in suppression by *tof1Δ* compared to *fob1Δ* is perhaps related to a more general role of Tof1 in replisome pausing throughout the genome, since Fob1 is thought to act exclusively at rDNA repeats. Altogether, our results show that Tof1 mediates fork pausing, rDNA instability and cellular toxicity in cells lacking Rrm3 helicase. Therefore, it is unlikely that Tof1 promotes fork pausing exclusively by regulating Rrm3 helicase but rather suggests a more direct involvement of Tof1-Csm3 in fork slowdown (**Fig. 1A**, model ‘2’), albeit through an unknown mechanism.

### Tof1-Csm3 complex interacts with Top1

Intrigued by the strong rDNA stabilizing effect of *tof1Δ* mutation (**Fig. 1E**), we sought to identify the factor(s) contributing to this stability and regulating replication fork pausing at Fob1-RFB. We carried out an unbiased forward genetic screen for mutants de-stabilizing the rDNA in either a wild type (WT) or *tof1Δ* background, using *ADE2* and *URA3* loss from the array as a read-out (the “cowcatcher” screen, Materials and Methods; **Fig. S2A**). Mutations in *RRM3*, *SIR2*, *HST3*, *CAC1*, *ORC1* and *PSF2* genes were recovered in the WT background but not in *tof1Δ.* One of the mutations we discovered specifically in the *tof1Δ* background was in the *TOP1* gene, which encodes topoisomerase I (**Fig. S2A**) – an enzyme required for both DNA replication and stability of rDNA repeats (Christman et al. 1988; Kim and Wang 1989b; Kim and Wang 1989a). The highly negative score of this *top1-G297D* mutation in Protein Variation Effect Analyzer (Choi et al. 2012) (PROVEAN: -7; cutoff = −2.5) implied a deleterious effect of this change on Top1 function. Indeed, complete deletion of the *TOP1* ORF led to a strong elevation of rDNA instability (**Fig. 2A**). In contrast to *rrm3Δ* and *rif1Δ* mutations however, the rDNA instability in *top1Δ* cells was not suppressed by *tof1Δ*, suggesting that Top1 and Tof1 may have overlapping roles. This and the fact that *TOF1* was originally identified in a yeast two-hybrid screen that employed a part of Top1 protein as a bait, as its name implies (‘TOpoisomerase I-interacting Factor 1’; (Park and Sternglanz 1999)), prompted us to focus further on this factor.

**Figure 2.**
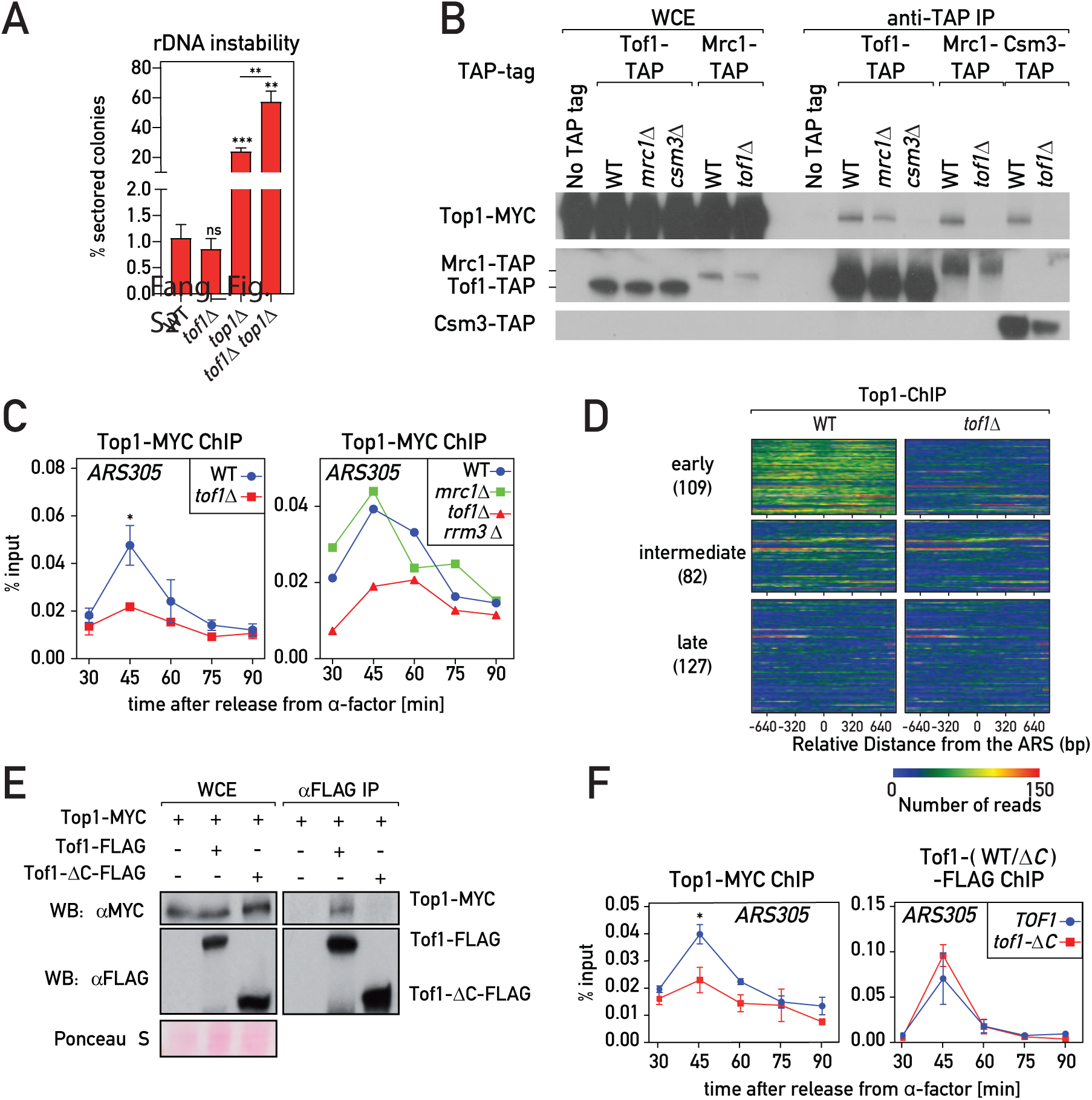
Tof1-Csm3-dependent recruitment of Top1 to the replisome (A) *tof1*Δ did not suppress *top1*Δ-induced rDNA instability, as measured with *ADE2* marker loss assay. (B) Top1 was co-immunoprecipitated with MTC complex in a Tof1- and Csm3-dependent but Mrc1-independent manner. (C-D) Chromatin DNA immunoprecipitated with Top1-MYC from cell cultures synchronously released into S phase from G1 (α-factor) arrest was: (C) subjected to qPCR on *ARS305*; (D) immunoprecipitated DNA from 45’ time point (early S phase) was Illumina sequenced, reads mapping to early, intermediate and late origins are shown as a heat map. (E-F) Tof1 lacking C-terminus (*tof1-*Δ*C* strains) did not co-immunoprecipitate Top1-MYC (E) and was defective in Top1-MYC association with *ARS305* during S phase (F). Here and on subsequent figures: *TOF1* = *TOF1-3xFLAG*; *tof1-*Δ*C* = *tof1*-Δ*981-1238-3xFLAG*. Values plotted and statistics as in Fig. 1. See also Fig. S2.

As mentioned above, Tof1-Csm3 is present in the cell nucleus within the MTC complex, together with Mrc1 (Bando et al. 2009). Using co-immunoprecipitation experiments, we observed that topoisomerase I was indeed recovered together with all the three components of the MTC complex (**Fig. 2B, S2B**). This interaction was detected only when whole cell extracts were treated with Benzonase nuclease, which degrades nucleic acids and liberates protein complexes from chromatin (De Piccoli et al. 2012) (**Fig. S2C**). Importantly, the MTC-Top1 interaction depended only on Tof1 and Csm3 proteins, but not Mrc1 (**Fig. 2B, S2B**), suggesting that Mrc1 interacts with Top1 indirectly through a Tof1-Csm3 sub-complex.

### Tof1-Csm3 promotes Top1 recruitment to the replisome

Since both Tof1-Csm3 and Top1 are components of the RPC (Gambus et al. 2006) we wondered whether the interaction of Tof1-Csm3 with Top1 occurs in the context of the replisome, which might explain how Top1 is recruited to the replication fork. To investigate this possibility, we conducted chromatin immunoprecipitation (ChIP) experiments to assess Top1 recruitment to origins of replication in cell cultures synchronously released into S phase from α-factor induced G1 arrest. We detected Top1 association with early origins (*ARS305* and *ARS607*) at the time of their activation (**Fig. 2C** and **S2D**) in accordance with a previous study (Bermejo et al. 2007). However, cells lacking Tof1 had much lower levels of Top1 recruitment to these (**Fig. 2C** and **S2D**). To confirm this result, we analyzed the genome wide binding of Top1 in early S phase and observed, as expected, that Top1 is enriched at replicating ARSs (**Fig. 2D**) and highly transcribed genes (**Fig. S2E**) that correspond to regions experiencing high helical tension. Remarkably, removing Tof1 abolished the Top1 signal at ARSs, whereas binding at promoters of highly transcribed genes was not affected. Furthermore, absence of the MTC complex member Mrc1 did not affect Top1 recruitment (**Fig. 2C** and **S2D**), which is in line with retention of the Tof1-Top1 interaction in *mrc1Δ* cells (**Fig. 2B** and **S2B**). Moreover, absence of the Rrm3 helicase did not restore the Top1 association with origins in *tof1Δ* cells (**Fig. 2C** and **S2D**).

The last 258 amino acid residues of the C-terminal part of Tof1 were reported to be sufficient for the two-hybrid interaction with Top1 (Park and Sternglanz 1999). Consistent with this part of Tof1 harboring a Top1-interacting domain, we observed a loss of Top1 co-immunoprecipitation and recruitment to origins in cells expressing a Tof1 protein lacking the last 258 aa (*tof1-ΔC = tof1-Δ^981-1238^-3xFLAG*) (**Fig. 2E-F** and **S2F-G**). Importantly, recruitment of WT Tof1 and the truncated Tof1-ΔC protein to origins was comparable (**Fig. 2F** and **S2F-G**). This suggests that Tof1 promotes Top1 association with origins by directly recruiting Top1 to the replisome.

### Top1 positively regulates replication fork pausing at RFBs

As it is not understood how Tof1-Csm3 slows down the replication fork at protein barriers, we wondered if their interactor Top1 is involved in this process. In order to assess this putative functional link between Tof1 and topoisomerase I, we evaluated replication pausing at RFBs in asynchronous cultures. Indeed, deletion of *TOP1* or dissociation of Top1 from the replisome by *tof1-ΔC* mutation led to a similar ca. 50% decrease in pausing at RFBs both in WT and *rrm3Δ* backgrounds, as detected by 2D and 1D gels at rRFB (**Fig. 3A** and **S3A-B**) or by Mcm4-MYC ChIP at rRFB and tRNA genes (**Fig. 3B** and **S3C**). Moreover, the fork pausing decrease in the double mutant *tof1-ΔC top1Δ* was comparable to that of single *tof1-ΔC* and *top1Δ* mutants (**Fig. 3B**), suggesting that the two factors could act in the same pausing pathway. Consistent with retention of Top1 recruitment to the FPC complex and to the replisome, *mrc1Δ* had no defect in pausing (**Fig. S3B**), as previously shown (Tourriere et al. 2005; Hodgson et al. 2007). Moreover, as the *mrc1Δ* mutation is known to decrease fork progression rates even more strongly than does *tof1Δ*, but has no effect on pausing (Tourriere et al. 2005; Hodgson et al. 2007), it seems unlikely that the decreased fork pausing in *tof1-ΔC* or *top1Δ* mutant could be an indirect consequence of any potential change in fork progression rates in these mutants.

**Figure 3.**
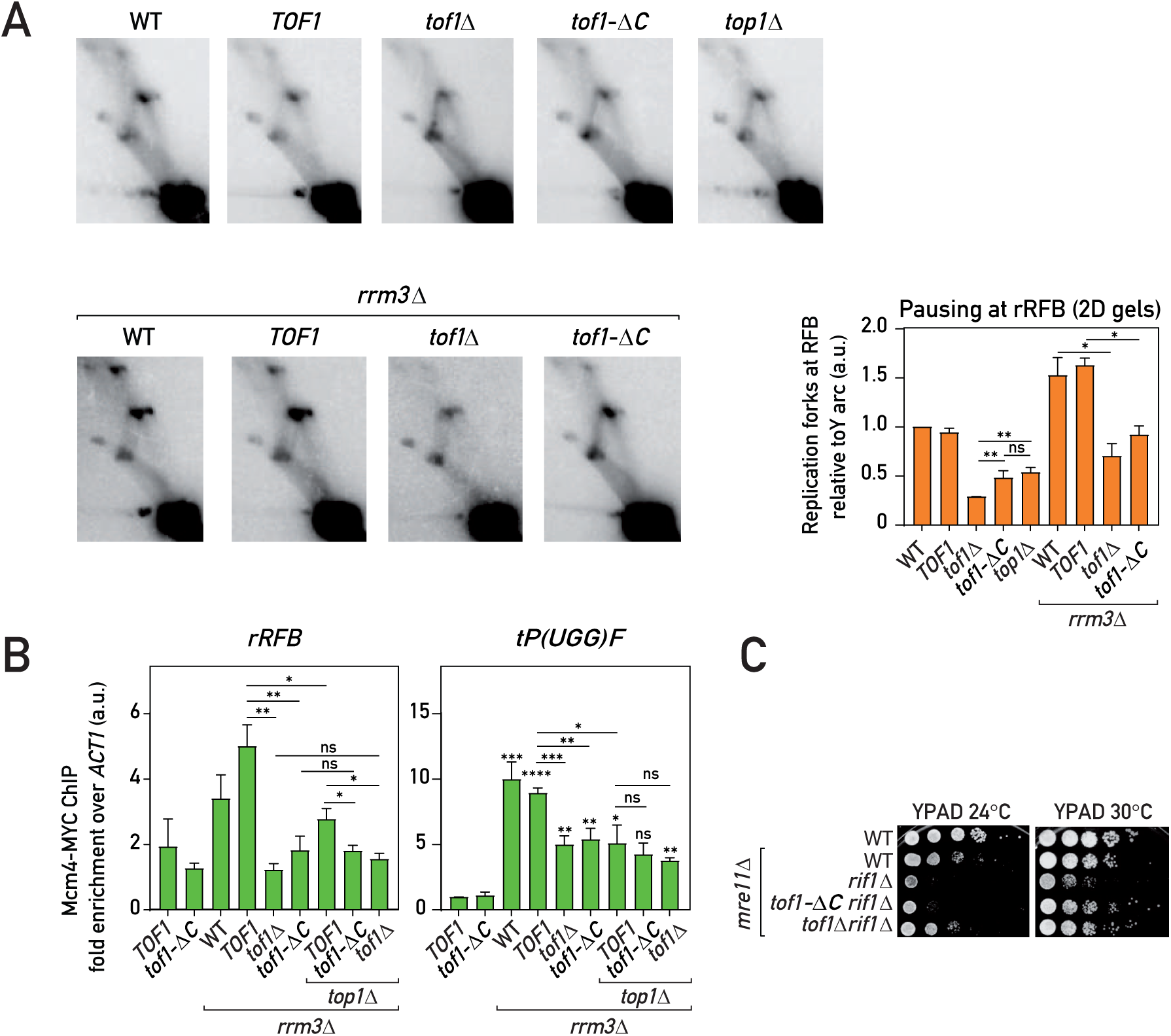
Tof1-C-dependent recruitment of Top1 to the replisome promotes fork pausing (A) Replication fork pausing at rRFB measured by 2D gels (as in Fig. 1B) in the strains of indicated genotypes: representative gel images and quantification (pausing in WT = 1, see Materials and Methods). (B) Replisome pausing at rRFB and a tRNA gene (*tP(UGG)F*) detected with Mcm4-MYC ChIP-qPCR in asynchronous cultures. (C) Tof1-ΔC is less toxic in *rif1*Δ *mre11*Δ cells than wild type Tof1. Values plotted and statistics as in Fig. 1. See also Fig. S3.

We had shown previously (Shyian et al. 2016) that *rif1Δ* leads to increased initiation at the rDNA ARS elements. One consequence of this is increased fork stalling and collapse at the rRFB, which leads to synthetic sickness in combination with *mre11Δ*. This synthetic growth defect is abolished by deletion of *FOB1*, confirming its connection to the rDNA fork block. As expected for a pausing defect, we found that *tof1-ΔC* partially alleviated *rif1Δ mre11Δ* synthetic sickness (**Fig. 3C**).

The fact that cells lacking Top1 completely or lacking the Top1-recruiting C-terminus of Tof1 still exhibit a pause signal significantly higher than cells lacking the whole of Tof1 protein (**Fig. 3A**) suggests that some other factor(s) are able to compensate for Top1 loss in a Tof1-dependent way and slow down the replisome in the absence of Top1.

### Top1 and Top2 redundantly promote fork pausing at Fob1-RFB

Top1 is believed to be the main replicative swivelase (Kim and Wang 1989a), but it is not essential for replication elongation and survival in budding yeast since Top2 is able to compensate for its absence (Kim and Wang 1989a; Bermejo et al. 2007). Consistent with this, we also detected Top2 in the immunoprecipitates of Tof1 and Csm3 proteins (**Fig. S4A**) and *tof1-ΔC* mutation only partially affected this association (**Fig. S4B**). We asked then whether Top2 could compensate for the loss of the Top1 in the replication fork pausing. Indeed, while inactivation of topoisomerase II at elevated temperature in a *top2-ts* strain or by auxin-induced degradation of the protein had only a little effect on pausing (**Fig. 4A-B** and **S4C-D**) doing so in cells lacking Top1 (*top1Δ)* or in cells with Top1 destabilized from the replisome (*tof1-ΔC*) led to a dramatic fork pausing loss phenotype similar to the one in *tof1Δ* cells (**Fig. 4A-B, S4C-D** and **S4F-G**).

**Figure 4.**
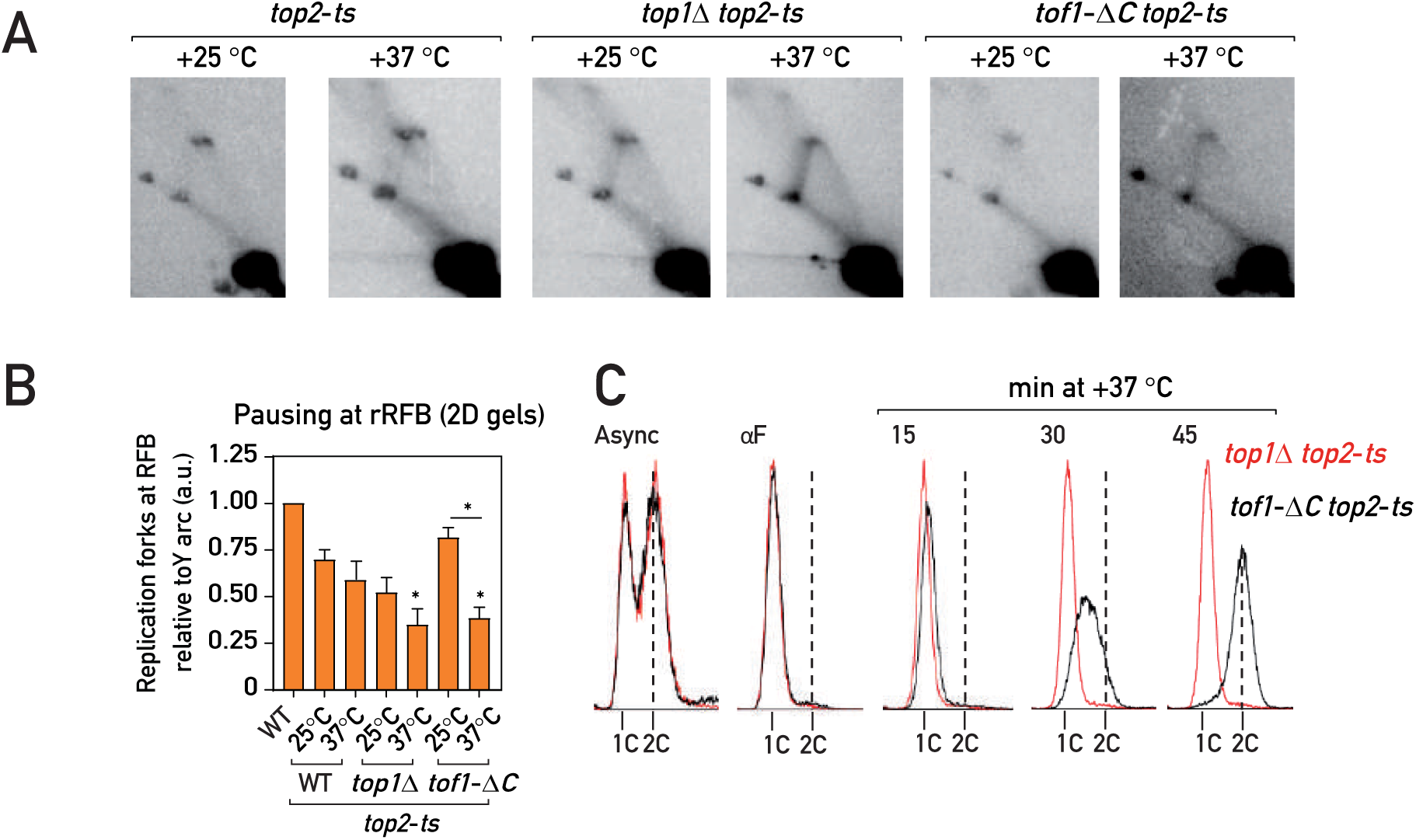
Top2 partially compensates for the fork pausing upon Top1 loss from the replisome (A-B) 2D agarose gel Southern blots (as in Fig. 1B): representative images (A) and quantification (B; pausing in WT = 1, see Materials and Methods) of replication intermediates in asynchronous cultures of the strains of indicated genotypes cultured continuously at +25 °C or transferred for 1 hour to +37 °C. (C) Flow cytometry DNA content profile of the *top1*Δ *top2-ts* (red) and *tof1-*Δ*C top2-ts* (black) strains upon release in S phase at +37°C from G1 (αF) arrest. Values plotted and statistics as in Fig. 1. Stars indicate P values for comparison with *top2-ts* strain at +25 °C. See also Fig. S4.

We observed a similar loss of fork slowdown when using different means to simultaneously deplete Top1 and Top2: temperature inactivation of Top2 in *top1Δ top2-ts* and *tof1-ΔC top2-ts* strains (**Fig. 4A-B** and **S4C**), degradation of both proteins (Top1-AID and Top2-AID) or degradation of Top2 in *top1Δ* and/or *tof1-ΔC* cells by the auxin-induced degron (Morawska and Ulrich 2013) system (**Fig. S4D-G**) and anchoring away (Haruki et al. 2008) of Top2 in a *top1Δ TOP2-FRB* background (**Fig. S4H**). Depletion of Top3 on its own or in combination with either Top1 or Top2 did not abolish the block (**Fig. S4D**), in accord with a recent study (Mundbjerg et al. 2015) and consistent with Top3 having a role in recombination but not replication (Pommier et al. 2016). Remarkably, replication intermediates in cells lacking both Top1 and Top2 had an appearance very similar to those of *tof1Δ* strains (**Fig. 3A, 4A**, and **S4C-H**), in which the loss of the pausing signal at the Fob1-RFB was accompanied by an increase in the intensity of the descending part (left half) of the Y arc. We speculate that the latter might be due to a head-on collision of the replication fork liberated from Fob1-RFB with the RNA polymerase I transcribing the adjacent rRNA gene. Thus, Top1 and Top2 proteins act in parallel to promote replication fork pausing at Fob1-RFB, and the replisome appears to be able to move past the Fob1-RFB in their absence.

Nevertheless, these data have to be interpreted with caution since it was reported that simultaneous inactivation of topoisomerase I and II leads to DNA damage checkpoint activation and to rapid replication cessation (Bermejo et al. 2007), which could in theory contribute to the observed fork pausing phenotypes. However, addressing the checkpoint issue, we found that degradation of both Top1 and Top2 in the checkpoint-deficient backgrounds *rad53-K227A* (kinase-dead Rad53) or *rad9Δ* abolished pausing to an extent similar to that in checkpoint-proficient cells (**Fig. S4H**), indicating that checkpoint activation is not necessary for the loss of replication fork slowdown. With regard to replication cessation, when released from G1 arrest into S phase at +37 °C, *top1Δ top2-ts* strains indeed failed to progress through S phase and arrested with close to 1C DNA content (**Fig. 4C**), consistent with previous findings (Kim and Wang 1989a; Bermejo et al. 2007). However, *tof1-ΔC top2-ts* and *tof1Δ top2-ts* cells rapidly progressed through the S phase in these conditions (similarly to *top2-ts* only cells (**Fig. 4C and S4E**)). Since both *top1Δ top2-ts* and *tof1-ΔC top2-ts* cells show a similar decrease of fork pausing at +37 °C (**Fig. 4A-B, S4C-D**), while only the former exhibits an S phase progression defect, we reasoned that the fork slowdown by Top1 and Top2 is not an indirect consequence of their genome-wide replication role but rather an *in cis* effect of these topoisomerases at the replisome, promoted by the Tof1-Csm3 complex. Moreover, it appears that Top1 (and perhaps Top2) anchoring at the replisome by Tof1 is not essential for the general S phase progression but is specifically important for fork pausing.

### tof1-ΔC is a separation of function mutation that leaves replication checkpoint roles intact

Since Tof1-Csm3 is an evolutionary conserved complex performing both fork pausing and replication checkpoint functions at the replisome (McFarlane et al. 2010), we wondered if the Tof1 C-terminus might be specifically involved in only the fork pausing role.

First, similar to the wild-type version, Tof1-ΔC protein appears to protect its partner Csm3 from degradation (**Fig. 5A**), an evolutionarily-conserved Tof1 function (Chou and Elledge 2006; Bando et al. 2009). Next, Tof1 positively regulates the DNA replication checkpoint (DRC) (Foss 2001), promoting survival of DNA damage response-deficient cells (*rad9Δ*) subjected to hydroxyurea-induced replication stress. Tof1-ΔC was still able to carry out this function (**Fig. 5B**), indicating that it is likely checkpoint-proficient. Accordingly, Tof1-ΔC supported DRC activation as measured by Rad53 phosphorylation in both *rad9Δ* and in WT cells (**Fig. 5C** and **5B**), while *tof1Δ rad9Δ* cells had a prominent defect in Rad53 phosphorylation similar to checkpoint defective *mec1Δ sml1Δ* cells, as expected (Foss 2001). Furthermore, Tof1 and Mrc1 appear to act in the same DRC pathway, as *tof1Δ* and *tof1Δ mrc1Δ* cells showed a similar defect in Rad53 phosphorylation under HU treatment (**Fig. 5D**), consistent with known role of the Tof1 in promoting Mrc1 association with replication forks (Bando et al. 2009).

**Figure 5.**
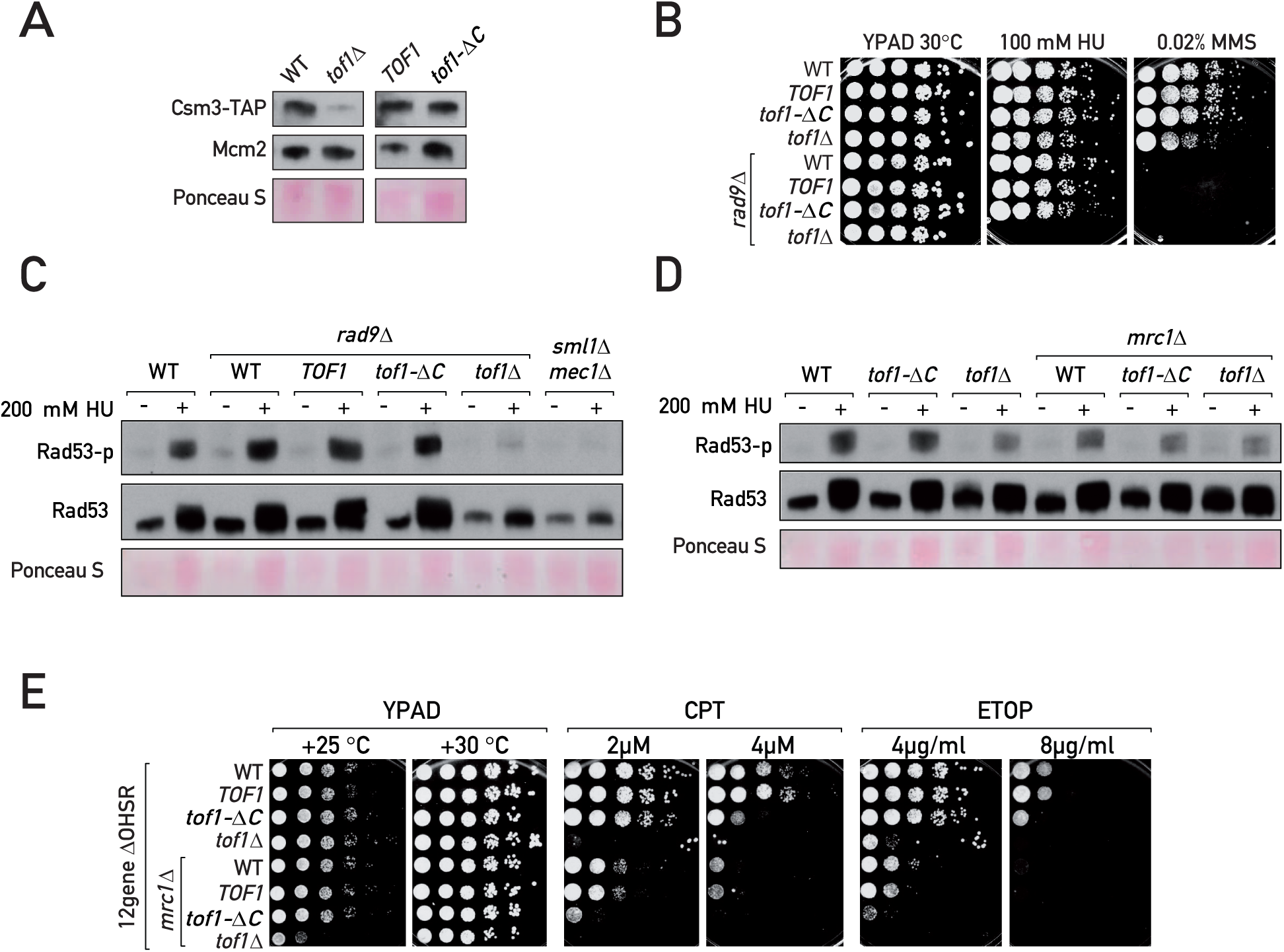
Fork pausing is a separable function of Tof1-Csm3 (A, C-D) Western blotting of TCA-extracted proteins. (A) In contrast to *tof1*Δ, *tof1-*Δ*C* cells do not degrade Csm3-TAP. (B, E) Serial dilution growth assays. (B) Tof1-ΔC supports viability of *rad9*Δ cells under hydroxyurea (HU) treatment. (C-D) Tof1-ΔC is proficient in DRC activation under HU treatment. (E) Mrc1 supports *tof1-*Δ*C* cells survival under topoisomerase-blocking damage. CPT – camptothecin; ETOP – etoposide; MMS – methyl methanesulfonate; 12geneΔ0HSR – multidrug sensitive yeast background. See also Fig. S5.

The loss of Tof1-Csm3 complex, but not Mrc1, confers strong sensitivity to the Top1-trapping agent camptothecin (CPT) (Redon et al. 2006; Reid et al. 2011) (**Fig. 5E** and **S5A-B**). We found in addition that cells lacking any of the MTC complex components display impaired growth in the presence of etoposide (ETOP) (**Fig. 5E** and **S5A-B**), a chemical blocking topoisomerase II, with *tof1Δ* and *csm3Δ* again having a greater effect than *mrc1Δ*. Importantly, *tof1Δ* and *csm3Δ* mutations impaired growth specifically in the presence of topoisomerase blocking agents but not upon DNA double-strand break induction by phleomycin or fork stalling and breakage by the alkylating agent MMS (**Fig. S5A-B**). Therefore, the Tof1-Csm3 complex appears to protect cells from blocked topoisomerases. We wondered whether this protection stems from the ability of Tof1-Csm3 to engage with Top1 and Top2. Surprisingly, *tof1-ΔC* mutant was still significantly resistant to CPT and ETOP (**Fig. 5E**). We reasoned that the higher sensitivity of *tof1Δ* to these agents in contrast to *tof1-ΔC* mutant could be due to the preservation of another function in the Tof1-ΔC protein and, since Tof1-ΔC is proficient in the Mrc1-dependent DRC (**Fig. 5C** and **4D**), speculated that this might also be related to a role shared with Mrc1. We therefore removed Mrc1 from the *tof1-ΔC* mutant cells and indeed observed an increase in CPT and ETOP sensitivity in the *tof1-ΔC mrc1Δ* double mutant, to an extent comparable to that of *tof1Δ* cells (**Fig. 5E**). Interestingly, *tof1Δ*, but not *tof1-ΔC*, grew slowly in combination with *mrc1Δ* (at 25^0^C; **Fig. 5E**) and in spore colonies, suggesting that Tof1-ΔC protein still performs an additional function important for growth in parallel to Mrc1. Thus, the Tof1-Csm3 complex appears to protect the cell from trapped topoisomerases by both interacting with them directly (through the C-terminus of Tof1, and perhaps other regions) and by acting together with Mrc1, likely by promoting the DRC and/or stabilizing forks at the topoisomerase-trapping sites (Strumberg et al. 2000).

## DISCUSSION

In summary, we showed that Tof1-Csm3 mediates replication fork pausing at proteinaceous RFBs through a pathway independent of Rrm3 helicase. Instead, Tof1-Csm3 complex interacts with topoisomerases I and II and mediates Top1 association with the replisome in normal S phase. Although we did not detect Top2 recruitment to replisomes in unchallenged cells with ChIP, alternative approaches should be used in the future to assess Top2 recruitment and its dependency on FPC. Top1 was previously identified as a part of the RPC (Gambus et al. 2006) and our report pinpoints the precise factor responsible for its engagement and suggests that eukaryotic cells do not rely exclusively on the DNA topology-mediated recruitment of topoisomerases to replicate chromosomes but rather have an association hub (Tof1-Csm3) to enrich them on the replisome. We imagine that this pathway could serve to prevent buildup of excess torsional stress in the vicinity of the replisome, by ensuring topoisomerase presence. This may avert uncontrolled escape of supercoils away from the fork by diffusion, supercoil ‘hopping’ (van Loenhout et al. 2012) or fork rotation (Schalbetter et al. 2015), possibilities that warrant further investigation.

Our findings indicate that either Top1 or Top2 is able to impose replication fork pausing at the Fob1-RFB, through a mechanism that we dub ‘sTOP’ (‘slowing down with TOPoisomerase I and II’) (**Fig. 6**). Indeed, it is assumed that in eukaryotes topoisomerase I and II act in front of the replication fork to unlink the parental DNA strands (Brill et al. 1987; Duguet 1997), while topoisomerase II acts also behind the fork to remove precatenanes. Either Top1 or Top2 is sufficient to assist in DNA replication elongation (Pommier et al. 2016), explaining why *TOP1* is not essential. The essential role of *TOP2* stems not from the replication elongation step, but from its crucial role in chromosome segregation during replication termination (Baxter and Diffley 2008). We imagine that the local increase in topoisomerase concentration/activity in the vicinity of replisome afforded by Tof1-Csm3 recruitment might assist general replication elongation by alleviating torsional stress. In cells lacking the Tof1-Csm3 complex topoisomerases would act more distributively but still ensure replisome progression, albeit perhaps less efficiently. We note that recruitment of an essential enzymatic function to the replisome by non-essential RPC factors is not unprecedented, since another RPC component, Ctf4, serves to recruit DNA polymerase α/primase (Simon et al. 2014) and Mrc1 stimulates the interaction of the leading strand DNA polymerase ε (Lou et al. 2008).

**Figure 6.**
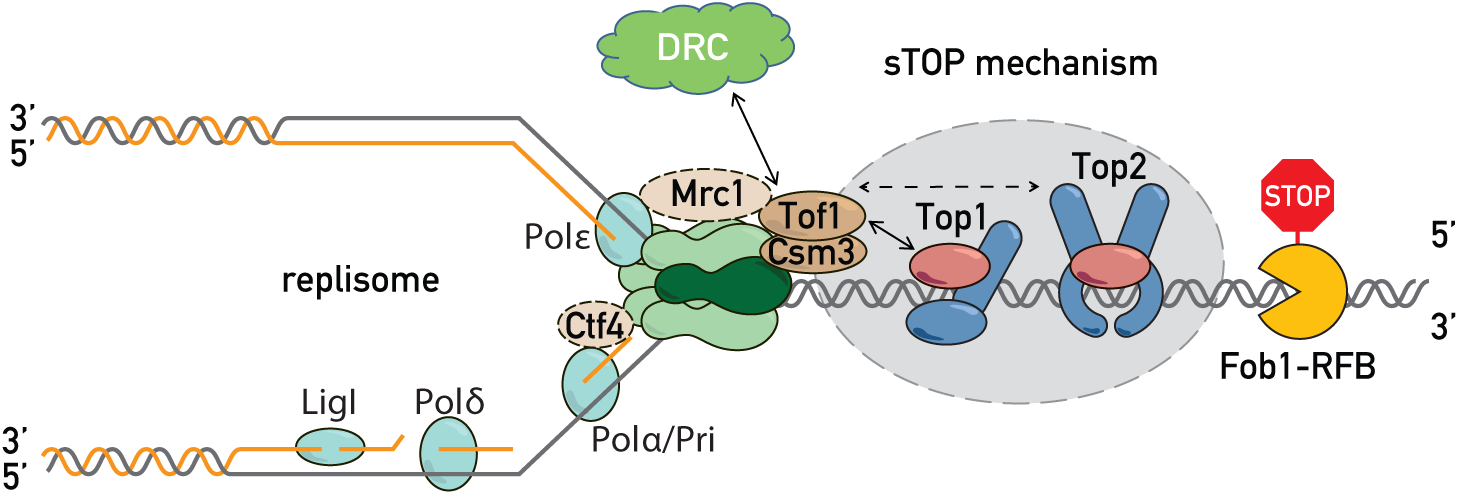
Replisome ‘sTOP’ model (‘slowing down with TOPoisomerases I-II’) Tof1-Csm3 promotes replication fork pausing at proteinaceous barriers via topoisomerase I (and II, dashed line with arrows), either by recruiting topoisomerases to the replisome or/and by recognizing topoisomarases bound in front of the fork. sTOP function of Tof1-Csm3 is distinct from its Mrc1-shared role in DRC (DNA replication checkpoint). See text for details.

In order to assist in DNA replication, the topoisomerase swivelase should be placed in front of the replication fork (Duguet 1997) – a setting where Top1 and Top2 might be the first replisome components to encounter obstacles. The slowing of the replication fork could be either a consequence of an inhibitory signal propagating from stalled topoisomerases through Tof1-Csm3 to the CMG helicase, or a result of topoisomerase activity itself. Consistent with first mode of action, it was reported that Tof1-Csm3 orthologues are able to inhibit the ATPase activity of MCM proteins *in vitro* (Cho et al. 2013). According to the second model, the absence of Top1 and Top2 at the replisome might promote bypass of barriers by increasing superhelical tension at the fork and simplifying blocking protein dissociation from DNA. In line with this possibility, bacterial *topA* mutants cause a loss of replication fork pausing at Tus/Ter sites likely by an increase in negative superhelicity, as this is suppressed by compensatory *gyrB* mutations (Valjavec-Gratian et al. 2005). Moreover, it was proposed that topoisomerase inhibition leads to nucleosome destabilization due to increased positive torsion ahead of transcribing RNA polymerase II (Teves and Henikoff 2014). It is thus tempting to speculate that by recruiting topoisomerases to the fork, Tof1-Csm3 precludes torsional stress buildup ahead of the replisome, helping to maintain integrity of chromatin (binding of both non-histone and histone proteins). Further studies, particularly with single-molecule approaches, will help to assess whether this is the case and elucidate the exact molecular details of how Top1 and Top2 promote the replication fork pausing at proteinaceous barriers and general fork progression.

Although topoisomerase would still be expected to assist DNA elongation by replisomes lacking the Tof1-Csm3 complex (since *tof1Δ* and *csm3Δ* cells are viable), the failure to recognize topoisomerases in front of the fork, and perhaps to duly pause until they dissociate from the template, might lead to replisome-topoisomerase collisions. We speculate that collision and replication run-off (Strumberg et al. 2000) with subsequent failure to properly activate checkpoint and repair the collapsed forks might explain the elevated sensitivity of *tof1Δ* and *csm3Δ* mutants to topoisomerase blocking conditions (**Fig. 5D, S5A-B**) (Redon et al. 2006; Reid et al. 2011). Accordingly, it was recently proposed that the Csm3 orthologue TIPIN may help to recognize topoisomerase I trapped by CPT and preclude replisome collision with it (Hosono et al. 2014).

This novel replication fork ‘sTOP’ mechanism offers a solution to an unresolved problem of how Tof1-Csm3 manages to recognize molecularly distinct RFBs: the Top1 and Top2 topological (or physical) interaction with RFBs might serve as a unifying common feature of different barriers. We also note that catalytically engaged Top1 is present at the Fob1-RFB (due to an interaction with Tof2 (Krawczyk et al. 2014)) throughout the cell cycle (Di Felice et al. 2005) and Top1 assists in progression of RNA polymerase II complexes (Teves and Henikoff 2014; Baranello et al. 2016). Therefore, an intriguing question would be whether Tof1-Csm3 could mediate recognition by the replisome of Top1 and Top2 present as a part of these and other chromatin complexes in the path of a replication fork.

Recently DNA replication elongation reactions (Yeeles et al. 2017) and fork pausing at Fob1 barriers were successfully reconstituted *in vitro* (Hizume et al. 2018), where Tof1-Csm3 supported high elongation rates and mediated pausing. It will be of interest to test whether these *in vitro* phenotypes of Tof1-Csm3 are mediated via recruitment of Top1 and Top2 to the replisome. Another fascinating question is whether the role of Tof1-Csm3 orthologues in other systems, such as replication pausing and imprinting control by Swi1-Swi3 at the *mat* locus in fission yeast (Dalgaard and Klar 2000), circadian clock regulation in metazoans (McFarlane et al. 2010), and survival in the face of replication stress (Bianco et al. 2019) are mediated via interactions with topoisomerases.

## Materials and Methods

### Yeast strains, genetics and growth conditions

Standard genetic methods for budding yeast strain construction and crossing were used (Shyian et al. 2016). Stains used in this study are listed in the **Table S1**. Genotoxic agent sensitivity was assessed in multidrug-sensitive yeast background (Chinen et al. 2011). For growth assays, saturated cultures of the respective genotypes were serially diluted (1:10) and spotted onto YPAD plates or YPAD plates supplemented with genotoxic agents. The plates were imaged following 2-4 days of incubation at 30°C or 25°C. *ADE2* marker loss assays were performed essentially as in (Shyian et al. 2016). Degradation of AID-tagged proteins (Morawska and Ulrich 2013) and cytoplasmic anchoring of the FRB-tagged proteins (Haruki et al. 2008) was achieved by addition of 1 mM IAA (Indole-3-acetic acid) or 1 mkg/mL RAPA (Rapamycin) for 60 minutes to the exponentially growing cultures. Heat inactivation of the Top2-ts protein was achieved by shifting exponentially growing yeast cultures from +25°C to +37°C for 60 minutes. For the cell cycle progression analysis in *top2-ts* background, the exponentially growing cells were arrested in G1 with αF treatment at +25°C during 2.5 hrs, transferred to +37°C for an additional 1 hr, washed 2x times with H2O, and released from the G1 arrest at +37°C in pronase-containing medium (Mattarocci et al. 2014).

### rDNA instability (ADE2 loss) assay

rDNA instability was assessed by the *ADE2* maker loss assay (Kaeberlein et al. 1999; Shyian et al. 2016). Saturated yeast cultures were diluted in water to around 400 cells per volume and plated onto YPD plates supplemented with 5 mg/ml adenine or onto SC plates (with or without 5-FOA). Plates were incubated at 30°C for 3 days, then at 4°C during 2 days and subsequently at 25°C for 1 day. The colonies were counted using ImageJ software Colony Counter plugin and the marker loss was plotted as the percentage of white colonies having red sectors to all the colonies except completely red colonies (where *ADE2* marker was lost in previous cell divisions).

### 1D and 2D gels and Southern blot

2D gels were performed essentially as in (Shyian et al. 2016) using BglII enzyme for genomic DNA digestion and Fob1-RFB Southern blot hybridization probe. The images were acquired with Typhoon FLA 9500 (GE Healthcare Life Sciences) and the intensity of signals quantified with ImageQuant TL 8.1 Software (GE Healthcare Life Sciences). The ratio of the signals at the rRFB spot to the remainder of Y arc of a given mutant was normalized to the respective ratio in WT present on the same 2D gel membrane and reported as ‘*Replication forks at RFB relative to Y arc’* value; this value in all the WT samples therefore equals 1. For 1D gels the first dimension gel was stained with EtBr, directly transferred to nylon membrane and probed with a radioactively labeled probe specific to Fob1-RFB site (Brewer and Fangman 1988; Kobayashi et al. 2004). The membranes were exposed to K-screens (Bio-Rad) for 6 hrs to 7 days before phosophorimaging.

### Chromatin Immunoprecipitation (ChIP)

Mcm4-13MYC, Cdc45-13MYC, Top1-13MYC anti-Myc and Tof1-3FLAG, Tof1-980aa-3FLAG anti-FLAG ChIP assays were performed essentially as in (Mattarocci et al. 2014). Mcm4-13MYC and Cdc45-13MYC ChIP experiments were done using asynchronously growing cultures. Where indicated, precipitated DNA was used to prepare sequencing libraries with TruSeq (illumine) and sequenced on iGE3 Genomics Platform of University of Geneva. FASTQ files were mapped to *S. cerevisiae* genome with Mapping tool of ‘HTSstation’ (David et al. 2014). Cell synchronization and flow cytometry assays were performed essentially as described in (Mattarocci et al. 2014).

### ‘Cowcatcher’ screen

Strains containing single copy of *ADE2* and *URA3* genes inserted into rDNA array were used for mutagenesis with EMS at 50% survival. EMS-treated cultures were split in 10 separate tubes, inoculated into SC-ADE-URA liquid medium and grown overnight (to counter-select mutations in *ADE2* and *URA3*). Then, aliquots were inoculated into YPAD and grown overnight to allow for marker loss from the rDNA. Dilutions were plated on 5-FOA plates (selection for *URA3* loss) and incubated as in *ADE2* loss assay above. After visual inspection, red sectored colonies from 5-FOA plates were manually selected and their white sectors were streaked sequentially 2 times onto SC plates. Of ca. 50’000 colonies from 5-FOA plates, 30 independent, reproducibly high-sectoring isolates were chosen. These were back-crossed, sporulated, dissected and assessed for segregation of the high sectoring phenotype. Isolates showing 2:2 segregation for sectoring (consistent with Mendelian mono-allelic mutations) were subjected to causative mutation identification using Pooled Linkage Analysis (as in (Birkeland et al. 2010; Lang et al. 2015)). Briefly, 20 spore colonies with a sectoring phenotype were pooled (+phenotype) and 20 white spore colonies were pooled (-phenotype) and their genomic DNA was isolated with a Qiagen genomic tip kit. Total genomic DNA of the two pools was submitted to iGE3 Genomics Platform of University of Geneva for fragmentation, library preparation and whole genome deep sequencing. The resulting FASTQ files were mapped to *S. cerevisiae* genome with Mapping tool of ‘HTSstation’ (David et al. 2014). The SNPs were identified with the SNP tool of ‘HTSstation’. The SNPs unique/over-represented in the plus-phenotype pool compared to the minus-phenotype pool were identified in Excel.

### Co-immunoprecipitation, SDS-PAGE and Western blot

Co-immunoprecipitation was performed as in (Gambus et al. 2006) and (De Piccoli et al. 2012). Briefly, 50 mL of exponentially growing cells at OD_600_ = 0.6 were pelleted, washed 2x times with cold H_2_O, suspended in 1 mL of Lysis Buffer (100 mM HEPES-KOH pH 7.9, 100 mM potassium acetate, 10 mM magnesium acetate, 10% glycerol, 0.1% NP-40, 2 mM EDTA, 2 mM glycerol 2-phosphate, and freshly added: 2 mM sodium fluoride, 1 mM DTT, 1 mM PMSF, Roche Protease Inhibitor Cocktail and PhosStop) and transferred into a cryotube with 500 μL of Zirconia/silica beads. The cells where homogenized in a Minibeadbeater at max power 2x times for 1.5 min with a 1 min interval. The lysed cells were recovered by centrifugation through a hole in the bottom of a cryotube and treated with 100 U of Benzonase (Millipore) for 40 min at +4°C with rotation. The whole-cell extract (WCE) was obtained as supernatant after centrifugation at 13000 rpm for 30 min at +4°C. 30 μL of IgG Sepharose beads pre-washed 4x times with Lysis Buffer were used for immunoprecipitation of the TAP-tagged proteins from the WCE during 2 hrs at +4°C with rotation. The beads were washed 3x times with Lysis Buffer at +4°C with rotation and boiled for 10 min with 50 uL of 2x Laemmli Buffer. The proteins were resolved on 8% iD PAGE GELS (Eurogentec), transferred onto nitrocellulose membrane (Amersham). The proteins were detected with anti-TAP (ThermoFisher), anti-MYC (Cell Singaling) or anti-FLAG (Sigma) antibodies. For Csm3-TAP protein level detection (Fig. 4A) and for Rad53 phosphorylation detection (Fig. 4C and 4D) total cellular proteins were isolated using TCA-Urea method (Mattarocci et al. 2014). Total and active auto-phosphorylated Rad53 were detected with Rad53 protein antibodies (Mab clone EL7) and (Mab clone F9) respectively provided by A. Pellicioli (University of Milan) (Fiorani et al. 2008).

### Statistical methods

Welch’s t test (two-tailed, unpaired t test with Welch’s correction) was used to assess statistical significance of differences in all the quantitative comparisons (*p<0.05; **p<0.01; ***p<0.001; ****p<0.0001). Mean values +/− SEM (standard error of the mean) are reported on graphs. GraphPad Prism 8 (GraphPad Software, Inc) was used to prepare the graphs and perform statistical comparisons.

## Acknowledgements

We thank Nataliia Serbyn for discussions, advice and assistance; Jessica Bruzzone for assistance with sequencing library preparation, Slawomir Kubik for advice on data analysis and all members of Shore Lab for critical comments. We thank iGE3 Genomics Platform of the University of Geneva for sequencing library preparation and high-throughput sequencing experiments. We thank Benoit Kornmann for the advice on the Pooled Linkage Analysis; Usui Takeo for sharing 12geneΔ0HSR background strains and Francoise Stutz for the BY deletion and DAmP strains. We thank Nicolas Roggli for expert graphical work; Pascal Damay for ordering and maintaining reagent stocks.

## Author contributions

MS conceived the project. MS, BA and DS designed all the experiments. MS, BA, AMZ, VI, GC and DD performed the experiments and analyzed the data. BA and VI contributed strains and reagents. MS wrote the manuscript, which was revised by MS and DS with input from BA.

---

**Figure S1.**
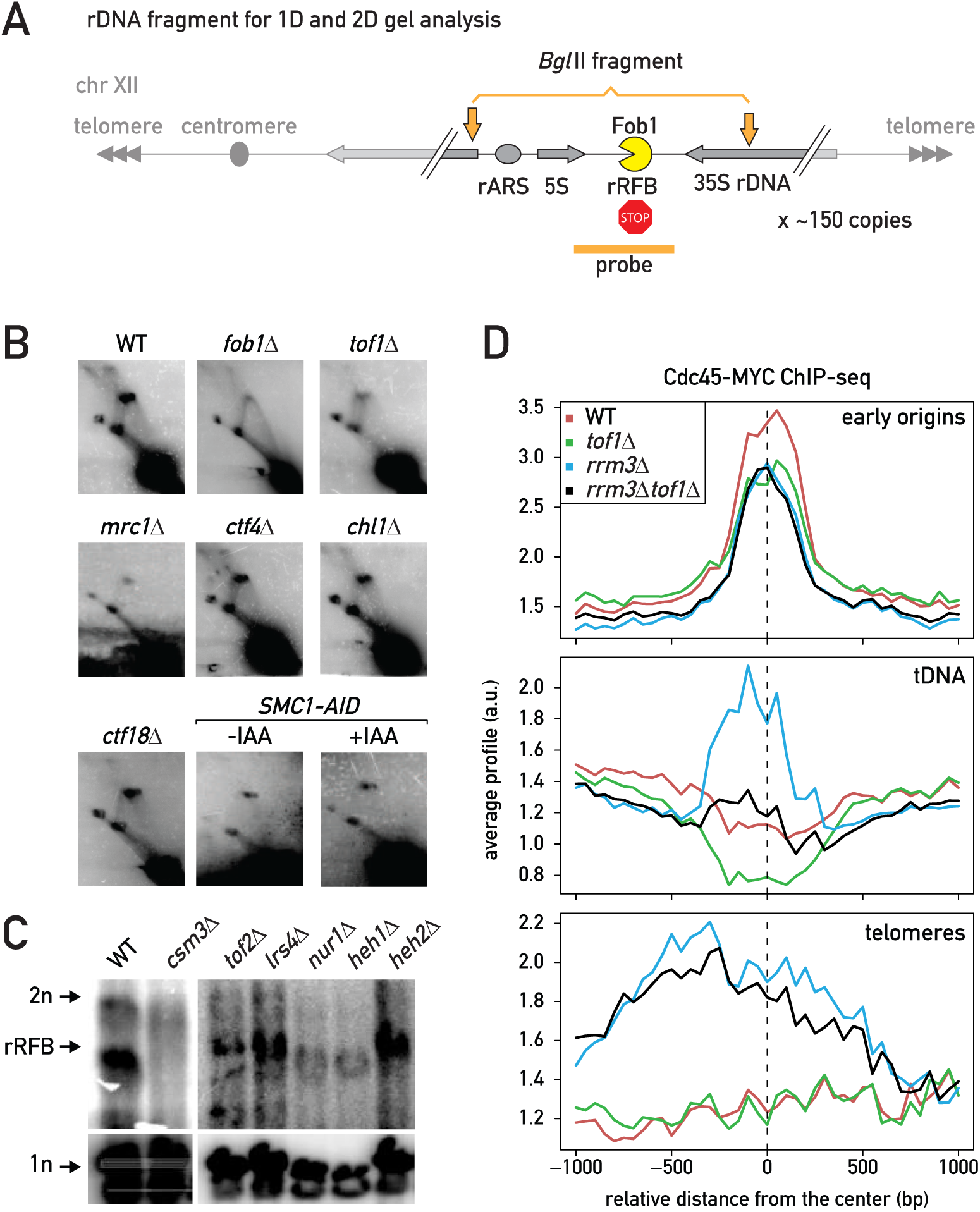
Related to Figure 1. Tof1-Csm3 complex functions independently of cohesion, peripheral anchoring and Rrm3 helicase (A) Diagram of the rDNA locus with the analyzed BglII fragment and location of the probe (rRFB) used for Southern blot hybridization of the 1D and 2D gels. (B) 2D gels for the estimation of fork pausing at rRFB in deletion mutants of the Replisome Progression Complex and sister chromatid cohesion establishment factors (*TOF1*, *MRC1*, *CTF4*, *CHL1*, and *CTF18*) and upon cohesin degradation with auxin (*SMC1-AID*). See Figure 1B for the diagram explaining DNA species on 2D gels. (C) 1D gels for the estimation of fork pausing at rRFB in deletion mutants of factors mediating peripheral anchoring of rDNA repeats to the nuclear envelope (cohibin and CLIP complexes). (D) Cdc45-MYC ChIP-seq in asynchronously growing cultures. Aggregation plots of the anti-MYC ChIP signal in Cdc45-MYC vs anti-MYC ChIP signal in WT not tagged control centered on early origins, tRNA genes and telomeres are shown.

**Figure S2.**
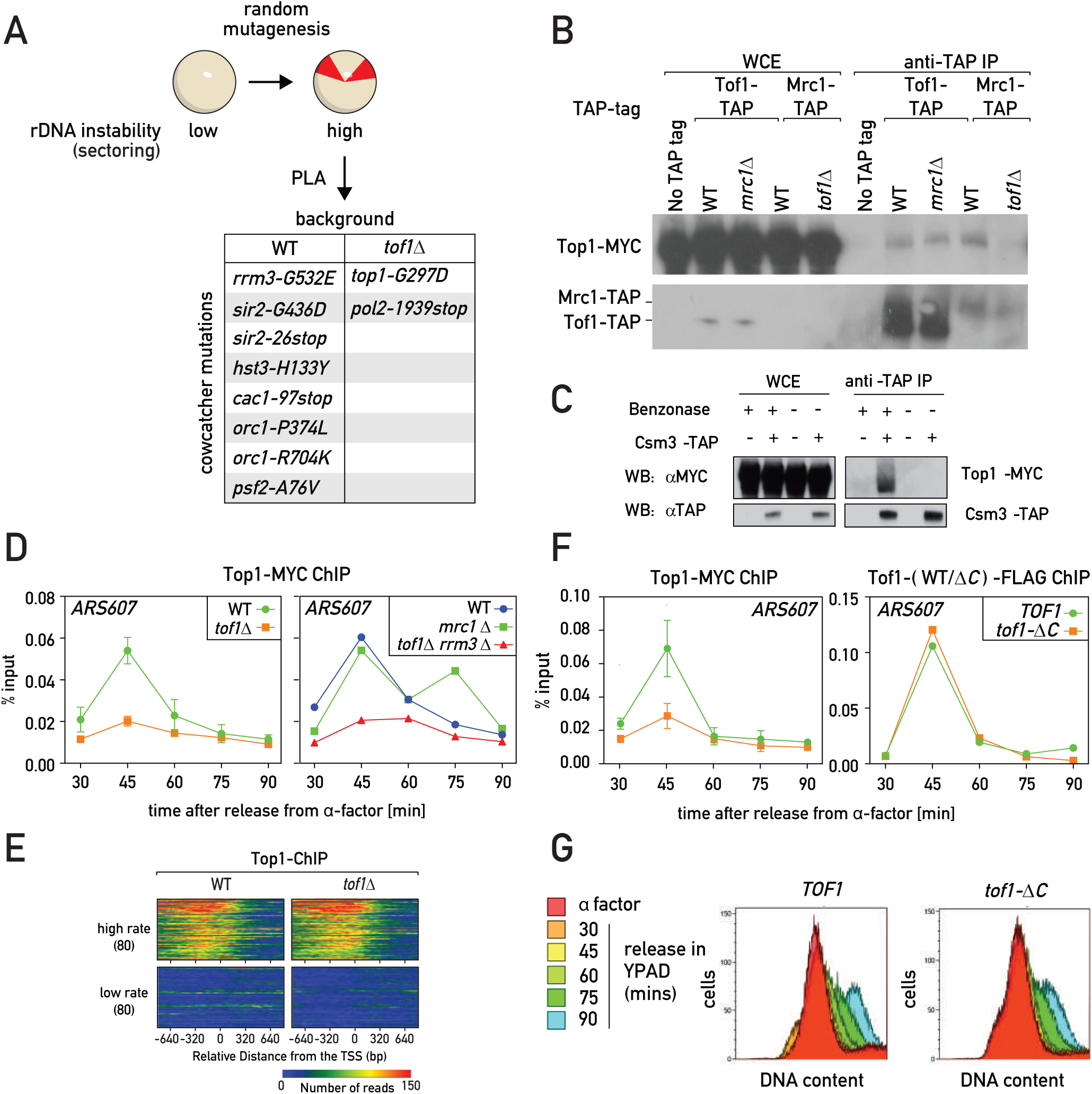
Related to Figure 2. Tof1-Csm3 recruits Top1 to the replisome (A) An outline of the “cowcatcher” screen and mutants identified by Pooled Linkage Analysis (PLA) as leading to elevated rDNA instability in WT and *tof1*Δ backgrounds. (B-C) Western blot detection of the Top1-MYC in Tof1-TAP and Mrc1-TAP (B) or Csm3-TAP (C) anti-TAP immunoprecipitates; DNA degradation by Benzonase was absolutely required to co-immunoprecipitate Top1 (C). (D-G) ChIP followed by qPCR at ARS607 or high throughput sequencing of Top1-MYC and Tof1- or Tof1-ΔC-FLAG in cell cultures synchronously released in S phase from G1 (α-factor) arrest: Top1-MYC ChIP-qPCR in the strains of designated genotypes (D); distribution of reads mapped to transcription start sites (TSS) is similar in WT and *tof1*Δ cells (E); Top1-MYC (left panel) or Tof1-FLAG and Tof1-ΔC-FLAG (right panel) ChIP-qPCR in *TOF1* and *tof1-*Δ*C* cells (F); flow cytometry profiles in *TOF1* WT and *tof1-*Δ*C* cells (for the experiment depicted on the Figure 2F and S2F) (G).

**Figure S3.**
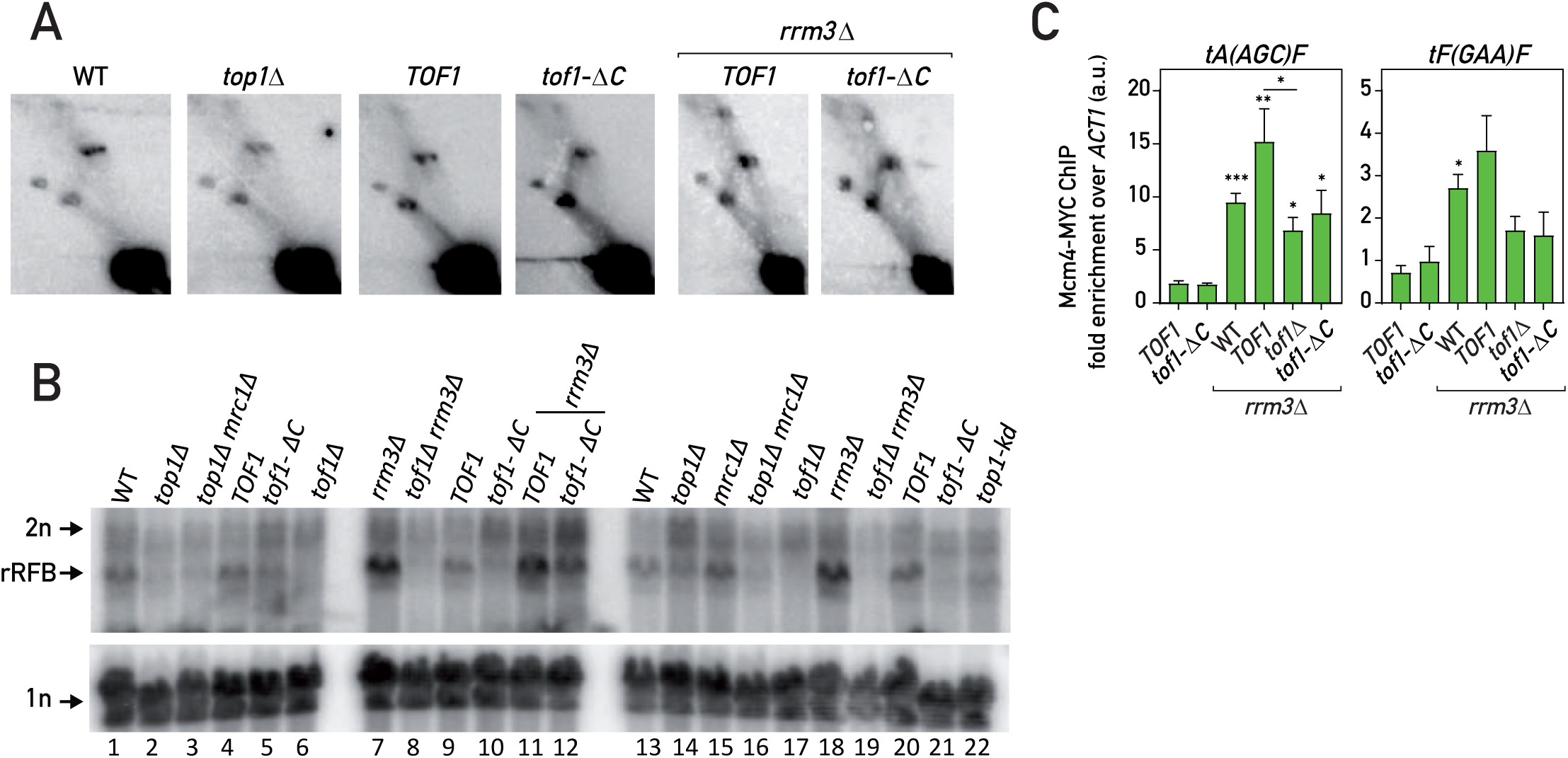
Related to Figure 3. Tof1-C with Top1 promote fork pausing (A-B) 2D gels (A) and 1D gels (B) of the *Bgl* II digested DNA isolated from asynchronous cell cultures and probed with rDNA rRFB probe. (C) Mcm4-MYC ChIP-qPCR in the asynchronous cultures of the strains of indicated genotypes for two tRNA genes. *top1-kd* – catalytically dead Top1 (*top1-Y727F*). Values plotted and statistics as in Figure 1.

**Figure S4.**
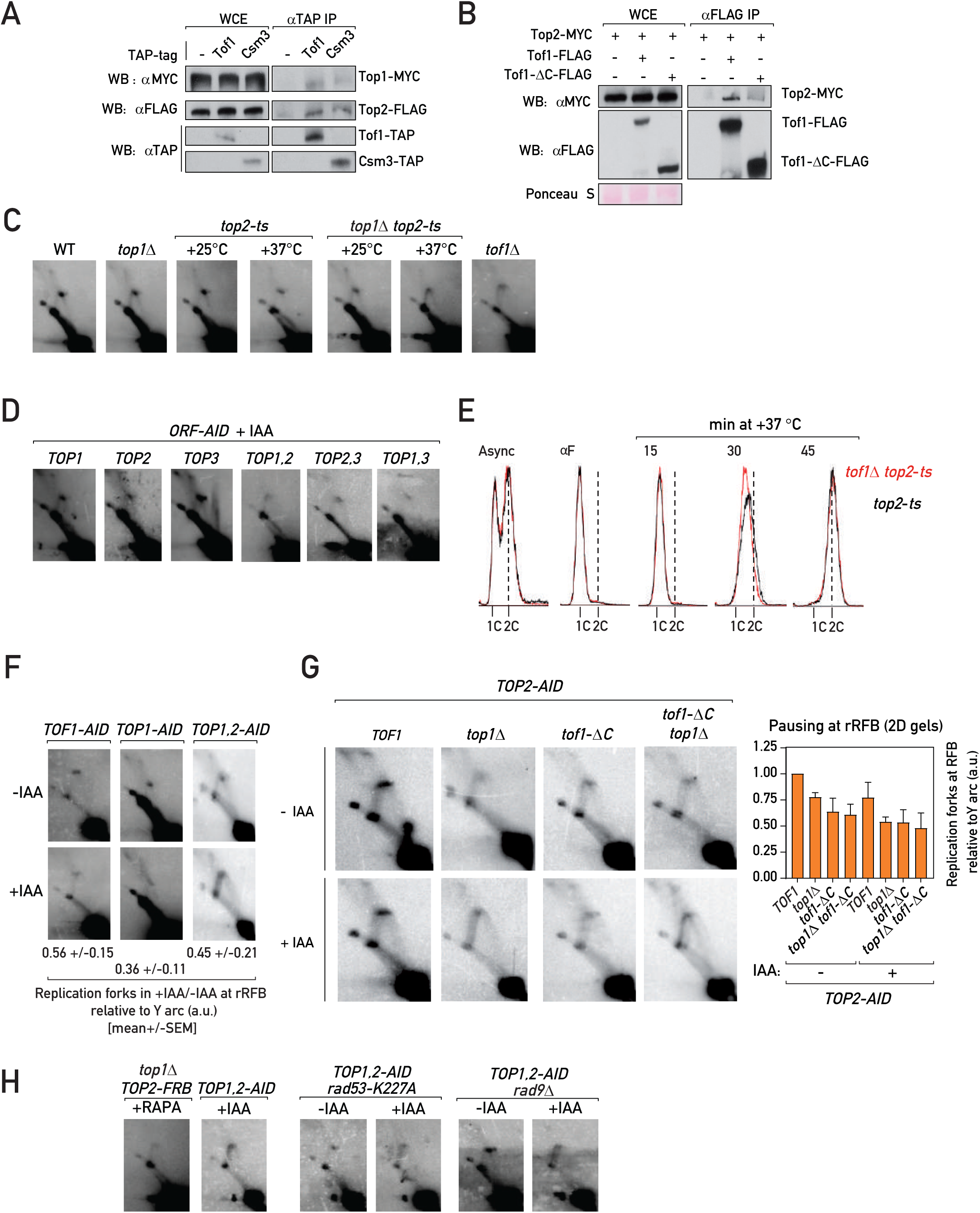
Related to Figure 4. Tof1-Csm3 engages Top1 and Top2 to pause the replisome (A-B) Co-immunoprecipitation experiments: Immunoprecipitates of Tof1 and Csm3 contained Top1 and Top2 (A), Top2 in the immunoprecipitates of Tof1-FLAG and Tof1-Δ C-FLAG (B). (C-D) 2D gels of the *Bgl* II digested DNA isolated from asynchronous cell cultures and probed with rDNA rRFB probe: DNA from asynchronous cultures of control strains (grown at +30 °C) or *top2-ts* strains (grown at +25 °C and shifted or not to +37 °C for 1 hour) (C); DNA from strains harboring AID-tagged topoisomerase genes *TOP1*, *TOP2* and *TOP3* from asynchronous cultures treated for 1 hour with 1 mM IAA (Indole-3-acetic acid) to degrade respective proteins (D). (E) Flow cytometry DNA content profile of the *tof1*Δ *top2-ts* (red) and *top2-ts* (black) strains upon release in S phase at +37 °C from G1 (αF) arrest. (F-H) 2D gels as in Figure S4D): upon Tof1, Top1 or Top1 and Top2 degradation (F); upon Top2 degradation (left panel – representative images; right panel – quantifications) (G); fork pausing in strains additionally harboring mutations of the DNA damage checkpoint genes (*rad53-K227A* and *rad9*Δ) (H). Values plotted and statistics as in Figure 1.

**Figure S5.**
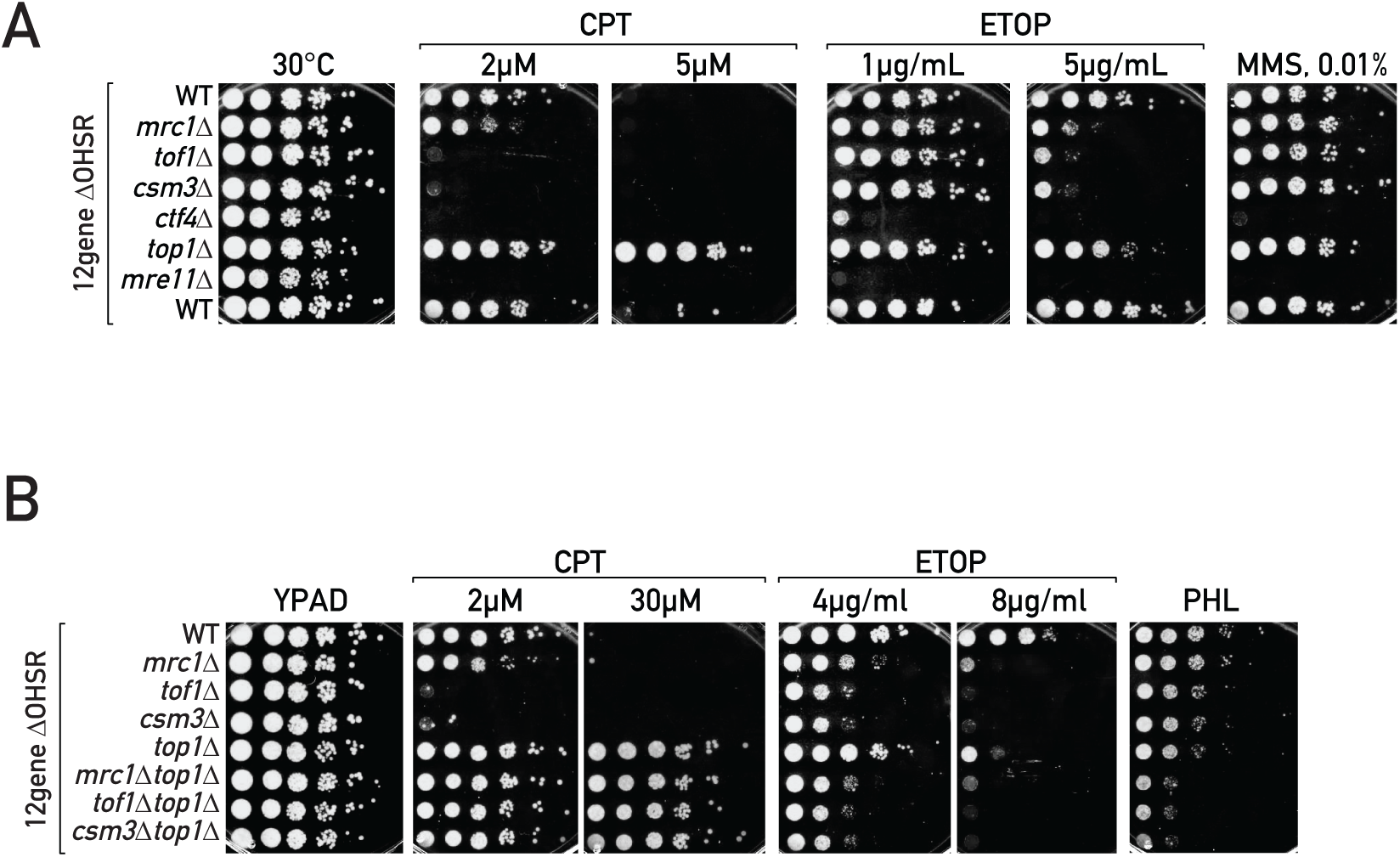
Related to Figure 5. Fork pausing complex protects cells from topoisomerase-blocking agents (A-B) Serial dilution growth assays: in mutants of replication fork progression complex (A) and in combinations of MTC genes deletion mutants and *top1*Δ. Genotoxic agents: CPT – camptothecin; ETOP – etoposde, MMS – methyl methanesulfonate (alkylating agent), PHL – phleomycin (used at 20 μg/mL; DSB-inducing agent).

**Table S1.**
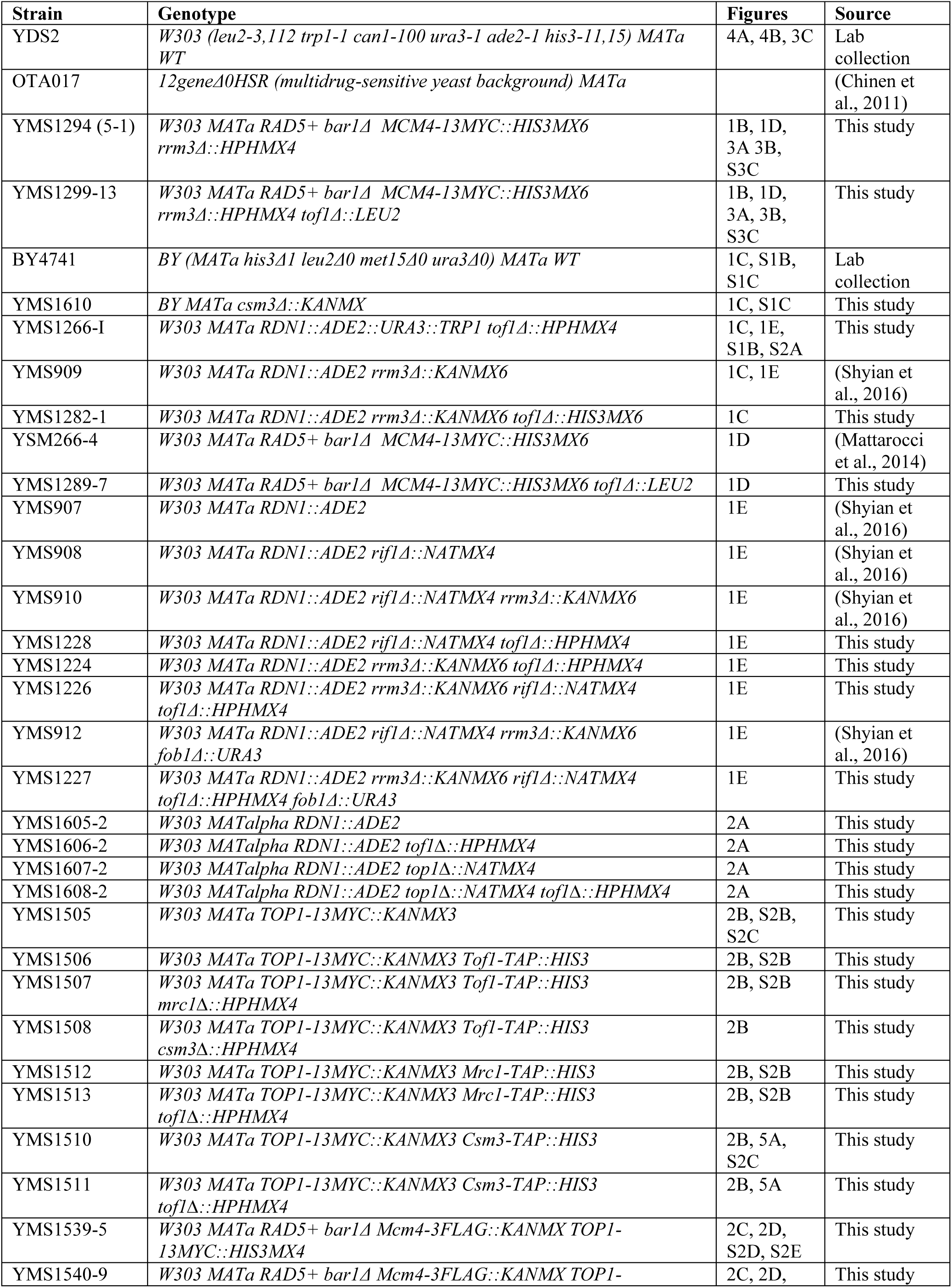
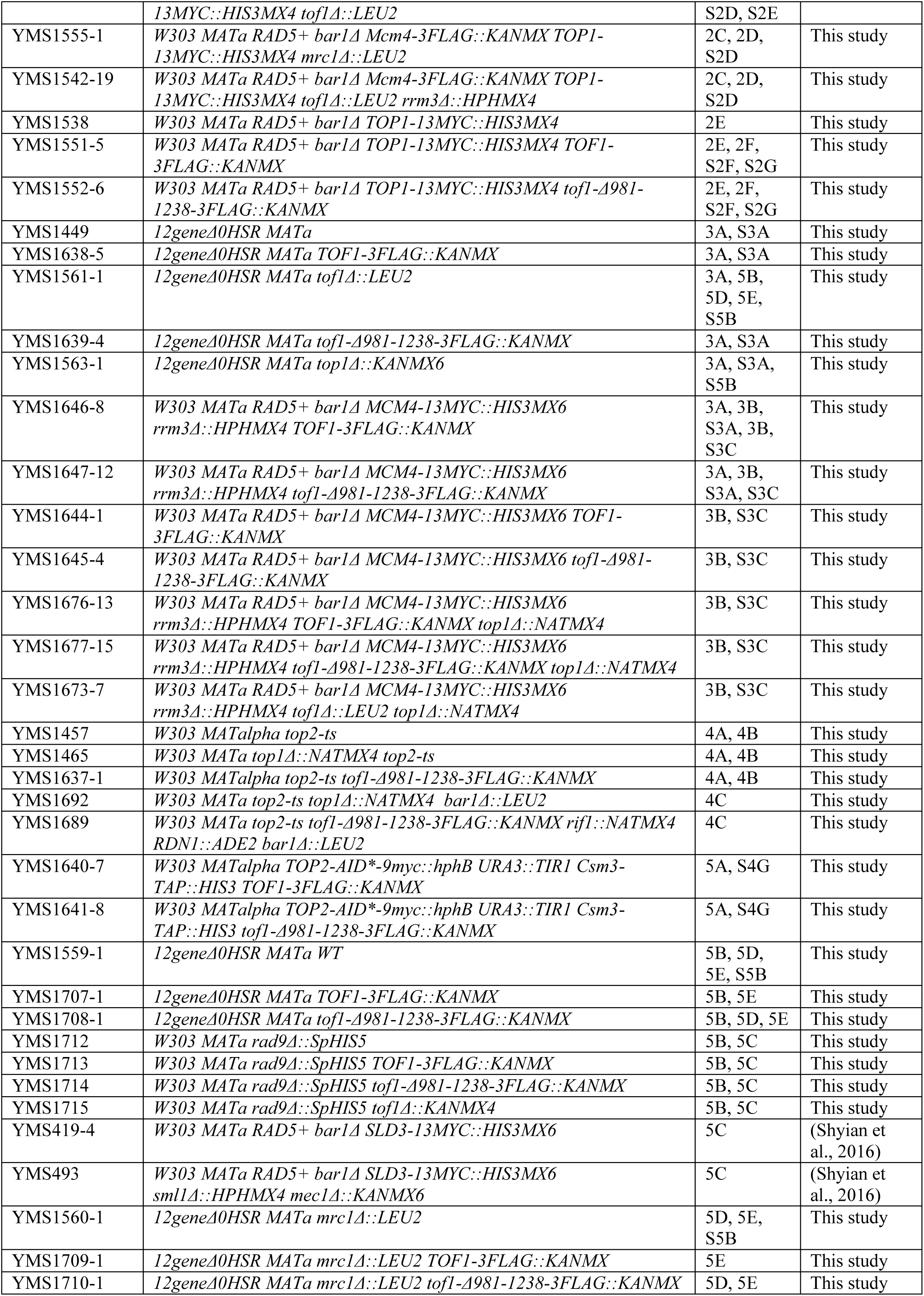
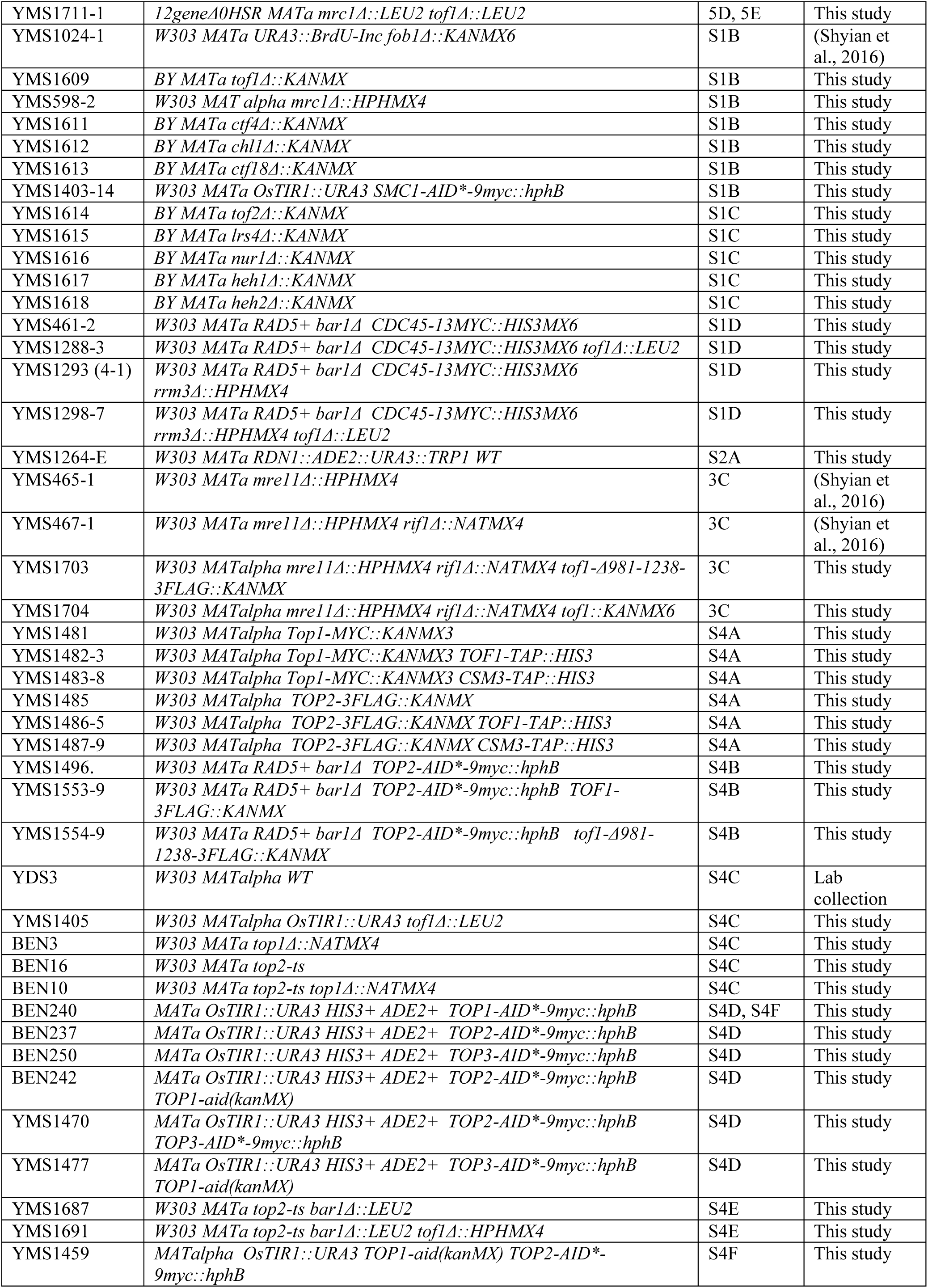
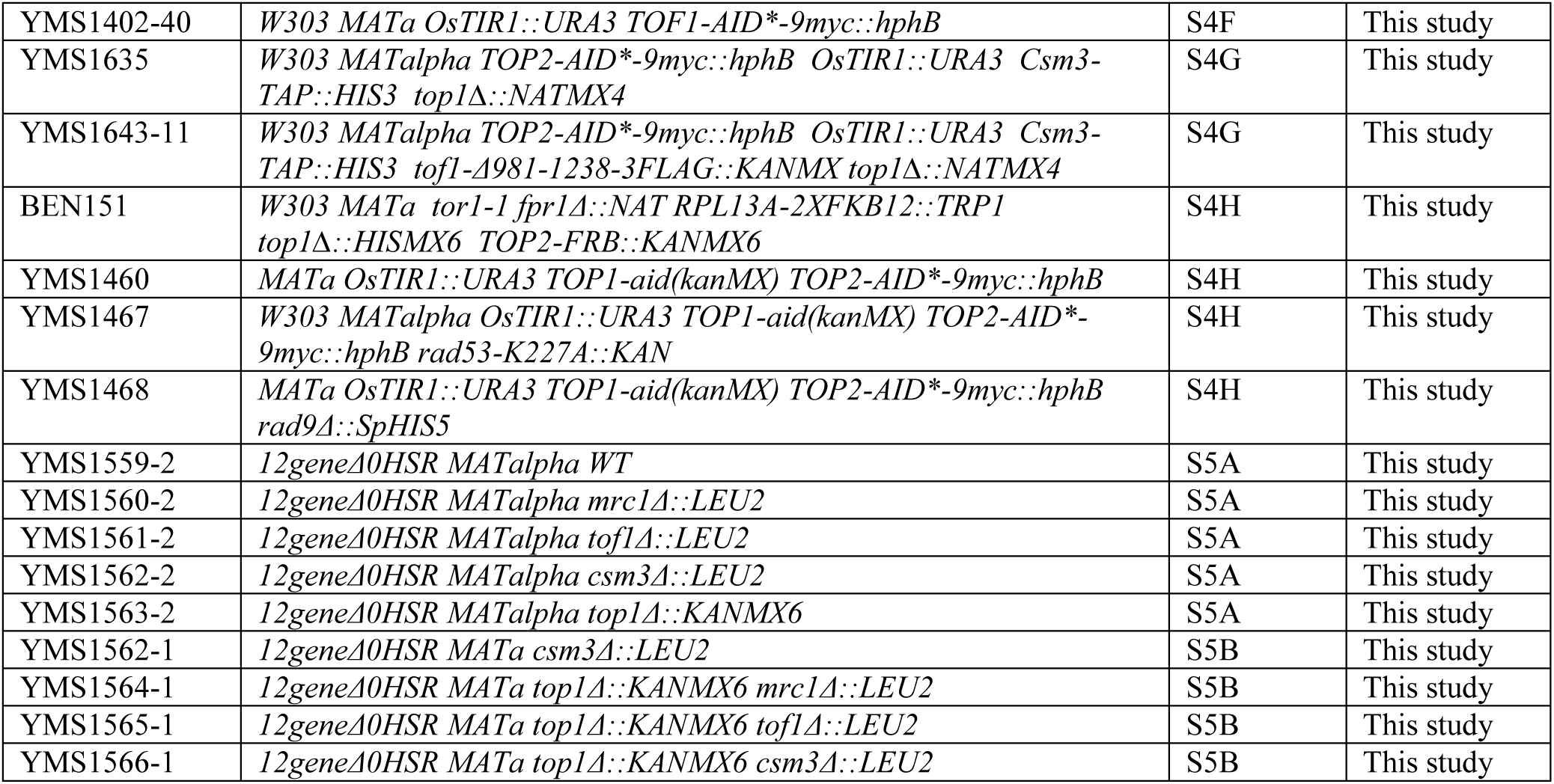
Yeast strains used in this study.

